## Supplementary Data for "Structural and molecular rationale for the diversification of resistance mediated by the Antibiotic_NAT family"

**Supplementary Fig. 1 – Sequence alignment of the Antibiotic_NAT family.**

Sequences are grouped and shaded according to phylogenetic reconstruction in Figure 1. Includes meta-AAC’s, AAC(3) enzymes from clinical isolates, and sequences identified through BLAST searches of NCBI. Darker color or shading of amino acids indicates higher conservation. Major and minor subdomains are indicated with solid and dashed black lines, respectively, above the sequence alignment.

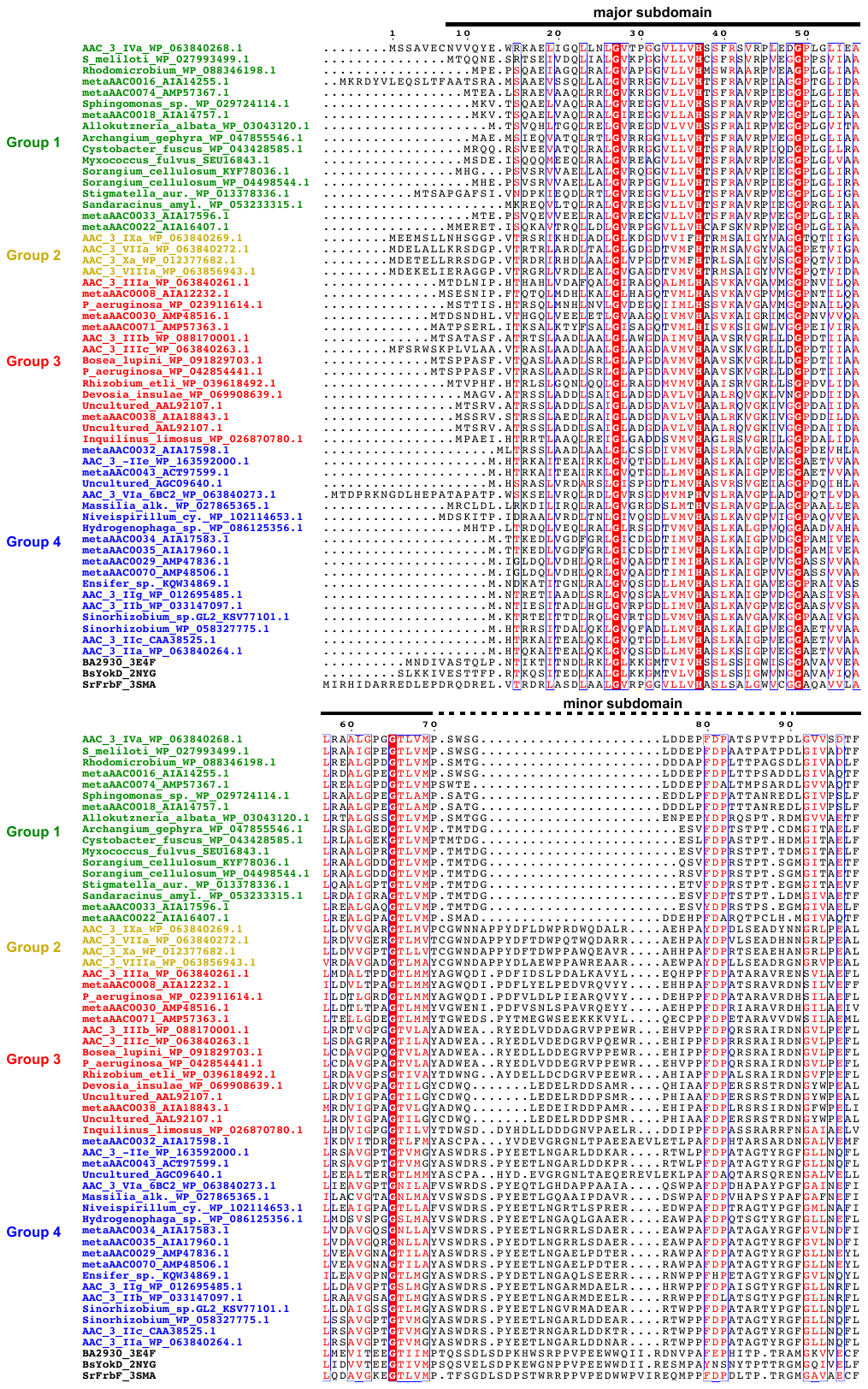

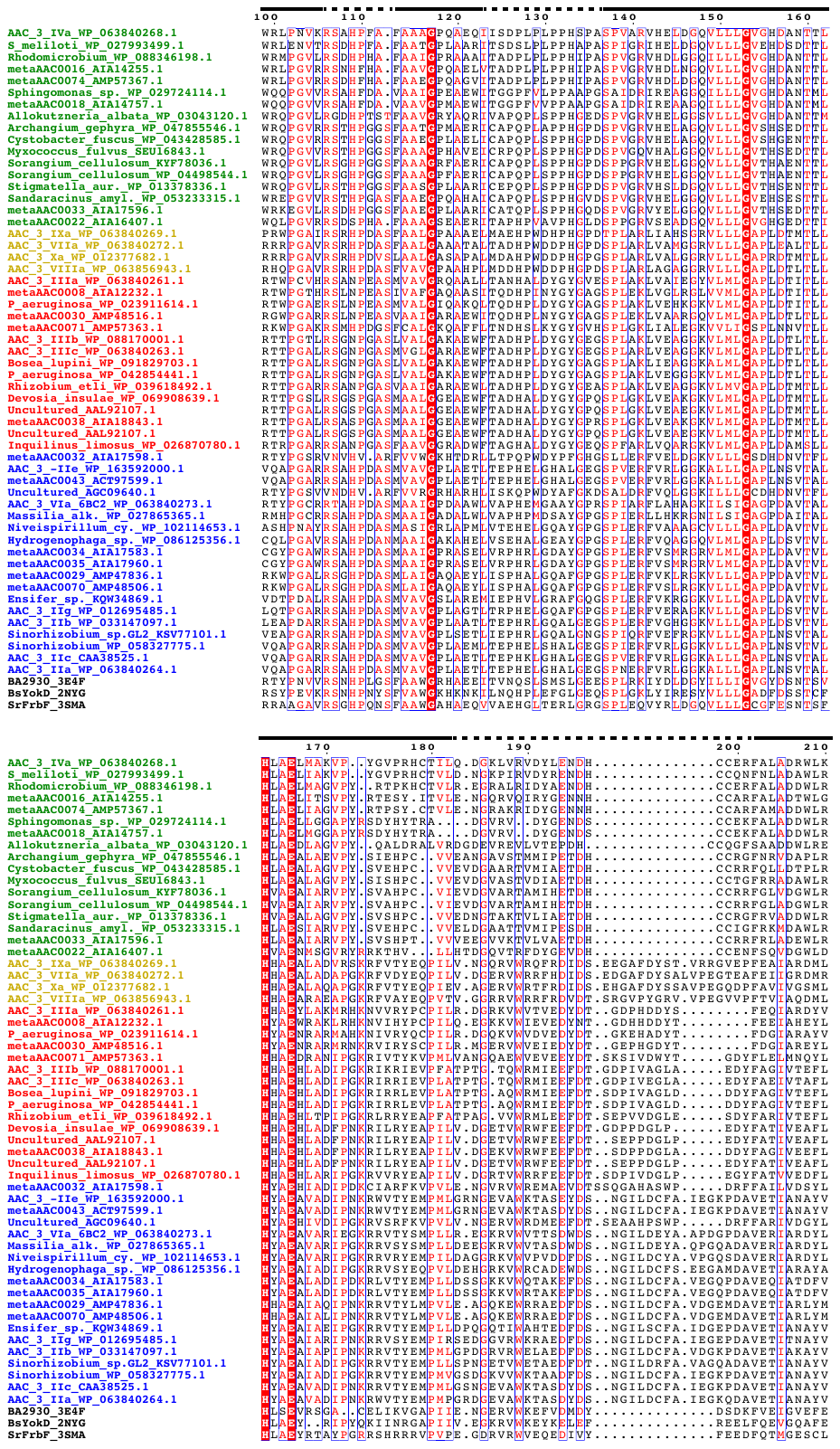

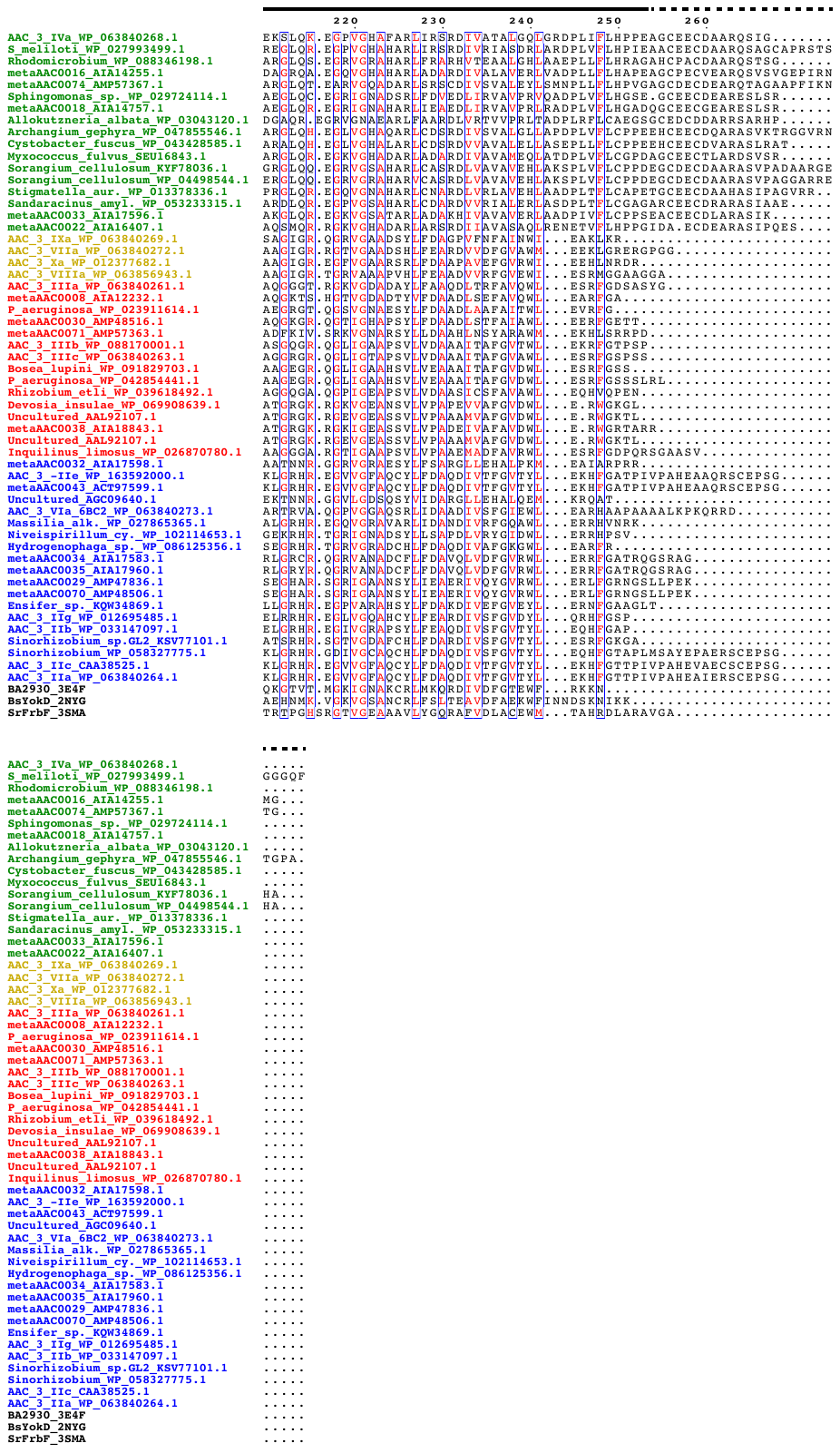

**Supplementary Fig. 2 – Structural analysis of Group 4 AAC(3)-IIb vs. AAC(3)-VIa.**

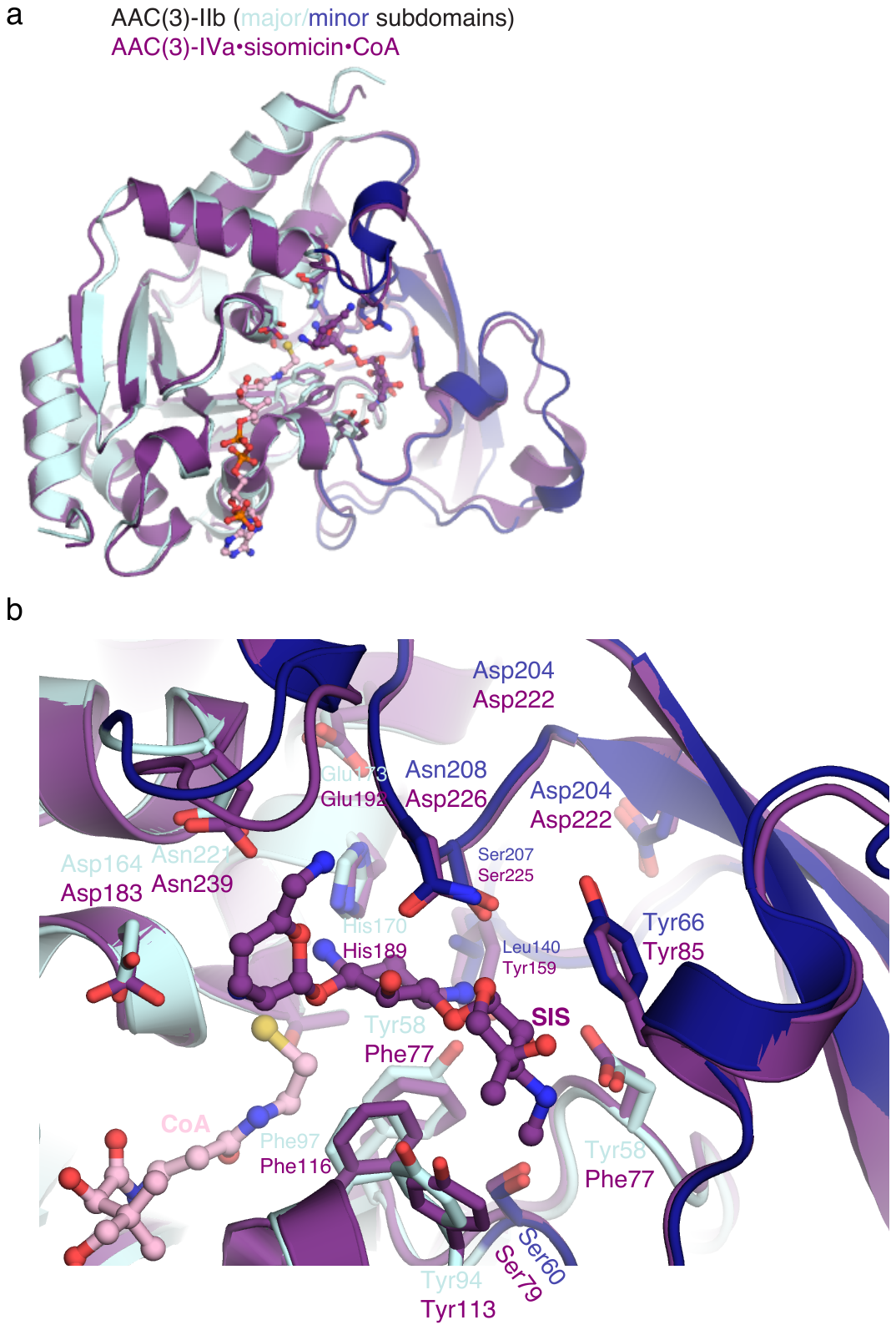

a) Superposition of overall structures. b) Zoom of active sites, sisomicin and CoA are shown in ball-and-stick representation.

**Supplementary Table 1. Aminoglycoside susceptibility of *E. coli* harboring Antibiotic_NAT genes**

*E. coli* BW25113 ∆*tolC*∆*bamB* expressing individual Antibiotic_NAT genes under the control of the P_bla_ promoter in vector pGDP3. Shown are MIC values in μg/mL grouped and coloured as in Figure 2.

|  |  | **Group 1** | | | | |
| --- | --- | --- | --- | --- | --- | --- |
|  | **Control** | **Meta-AAC0016** | **Meta-AAC0018** | **HMB0022** | **HMB00033** | **AAC(3)-IVa** |
| **APR** | 4 | ≥ 256 | ≥ 256 | > 256 | > 256 | ≥ 512 |
| **TOB** | 0.5 | > 256 | ≥ 256 | > 256 | > 256 | ≥ 64 |
| **GEN** | 0.25-0.5 | ≥ 256 | 128-256 | 64 | ≥ 256 | ≥ 512 |
| **KAN** | 2 | 128-256 | 64-128 | 8 | 32 | 64 |
| **AMI** | 1-2 | 1-2 | 1 | 1-2 | 1 | 1 |
| **NEO** | 1 | 64 | 128 | 8-16 | 64-128 | 64-128 |
| **PAR** | 2 | ≥ 256 | 128 | 16 | 64-128 | 128-256 |

|  |  | **Group 2** | | | |
| --- | --- | --- | --- | --- | --- |
|  |  | **AAC(3)-VIIa** | **AAC(3)-VIIIa** | **AAC(3)-IXa** | **AAC(3)-Xa** |
| **APR** |  | ND | ND | 4 | 2-4 |
| **TOB** |  | ND | ND | ≤ 0.5 | 16-32 |
| **GEN** |  | ND | ND | ≤ 0.5 | 8 |
| **KAN** |  | ND | ND | 2 | 64-128 |
| **AMI** |  | ND | ND | 1 | 1 |
| **NEO** |  | ND | ND | 1 | 1 |
| **PAR** |  | ND | ND | 2 | 4 |

|  |  | **Group 3** | | | | | |  |
| --- | --- | --- | --- | --- | --- | --- | --- | --- |
|  |  | **Meta-AAC0008** | **Meta-AAC0030** | **Meta-AAC0038** | **Meta-AAC0071** | **AAC(3)-IIIa** | **AAC(3)-IIIc** | **AAC(3)-IIIb** |
| **APR** |  | 16-32 | 16 | 32-64 | 4-8 | 8-16 | ND | 8 |
| **TOB** |  | > 256 | > 256 | > 256 | ≥ 256 | > 256 | ND | > 256 |
| **GEN** |  | > 256 | > 256 | > 256 | ≥ 256 | > 256 | ND | > 256 |
| **KAN** |  | > 256 | > 256 | > 256 | ≥ 256 | > 256 | ND | > 256 |
| **AMI** |  | 1 | 1-2 | 1-2 | 1-2 | 1 | ND | 0.5 – 1 |
| **NEO** |  | > 256 | 64-128 | 256 | 1 | 128-256 | ND | 256 |
| **PAR** |  | > 256 | > 256 | > 256 | 4 | ≥ 256 | ND | ≥ 256 |

|  |  |  | | | **Group 4** | | | | | | |  |
| --- | --- | --- | --- | --- | --- | --- | --- | --- | --- | --- | --- | --- |
|  |  | **Meta-AAC0029** | **Meta-AAC0032** | **Meta-AAC0035** | | **Meta-AAC0043** | **Meta-AAC0070** | **AAC(3)-IIa** | **AAC(3)-IIb** | **AAC(3)-IIc** | **AAC(3)-VIa** | **Meta-AAC0034** |
| **APR** |  | 4 | 2-4 | 8 | | 4 | ND | 8 | 16-32 | 2 | 4 | 8 |
| **TOB** |  | 64 | 4-8 | > 256 | | 32-64 | ND | ≥64 | > 256 | 64 | 8 | > 256 |
| **GEN** |  | > 64 | 64 | > 256 | | ≥ 256 | ND | ≥64 | > 256 | > 256 | 256 | > 64 |
| **KAN** |  | 16 | 16-32 | ≥ 256 | | 16 | ND | ≥64 | > 256 | 32 | 16 | > 256 |
| **AMI** |  | 0.5-1 | 1-2 | 1 | | 1-2 | ND | 2 | 1 | 0.5-1 | 1-2 | 1-2 |
| **NEO** |  | ≤ 1 | 1 | 1 | | 1 | ND | 1 | 0.5-1 | 0.5-1 | ≤ 1 | ≤ 1 |
| **PAR** |  | 1 | 2 | 2 | | 2-4 | ND | 2-4 | 1-2 | 0.5-1 | 2 | 1 |

ND = No data.

**Supplementary Table 2. X-ray crystallographic statistics.**

| **Structure** | **Meta-AAC0038^H29A^ apoenzyme** | **Meta-AAC0038^H168A^•AcCoA** | **Meta-AAC0038^H168A^•CoA** | **Meta-AAC0038^H168A^•apramycin•CoA** |
| --- | --- | --- | --- | --- |
| **PDB code** | 6MMZ | 6MN0 | 5HT0 | 7KES |
| **Data collection** |  |  |  |  |
| Space group | C2 | C2 | C2 | P3_1_2_1_ |
| Unit cell  *a*, *b, c* (Å)  α, β, γ, (°) | 105.8, 158.1, 143.4  90, 94.9, 90 | 108.1, 159.6, 143.3  90, 94.6, 90 | 107.02, 159.50, 146.22  90, 94.7, 90 | 127.77, 127.77, 94.65  90, 90, 120 |
| Resolution, Å | 25.00 – 3.30 | 25.00 – 2.40 | 25.0 – 2.75 | 30.0 – 2.36 |
| R*_merge_^a^*  R*_pim_* | 0.268 (0.743)  0.142 (0.395) | 0.094 (0.372)  0.062 (0.249) | 0.074 (0.440)  0.085 (0.251) | 0.091 (1.427)  0.031 (0.505) |
| CC_1/2_ | 0.809* | 0.949 | 0.968 | 0.601 |
| *I* / σ(*I)* | 6.3 (2.3) | 10.75 (2.09) | 17.76 (3.19) | 21.87 (1.0) |
| Completeness, % | 99.4 (99.9) | 99.9 (100) | 96.7 (90.4) | 100 (100) |
| Redundancy | 4.6 (4.6) | 3.3 (3.2) | 4.0 (3.7) | 9.9 (8.8) |
| **Refinement** |  |  |  |  |
| Resolution, Å | 19.75 – 3.30 | 24.93 – 2.39 | 24.97 – 2.75 | 29.19 – 2.36 |
| No. unique reflections:  working, test | 35122, 1646 | 94715, 1996 | 60061, 2021 | 36879, 1846 |
| *R*-factor/free *R­*-factor^b^ | 20.4/26.1 (29.6/38.9) | 17.8/20.8 (23.1/28.9) | 20.4/23.3 (31.3/30.8) | 19.1/22.8 (29.6/34.7) |
| No. refined atoms, molecules  Protein Aminoglycoside  Acetyl-CoA/CoA  Solvent  Water | 11977, 6  N/A  N/A  104  104 | 12033, 6  N/A  306, 6  236  1706 | 12016, 6  N/A  288, 6  105  343 | 3992, 2  73, 2  96, 2  25  170 |
| *B*-factors  Protein  Aminoglycoside  Acetyl-CoA/CoA  Solvent  Water | 59.2  N/A  N/A  96.2  20.4 | 32.9  N/A  33.1  71.4  43.4 | 54.1  N/A  52.9  108.2  47.7 | 70.9  129.0  61.9  100.5  64.2 |
| r.m.s.d.  Bond lengths, Å  Bond angles, ° | 0.002  0.552 | 0.005  1.770 | 0.014  1.827 | 0.005  1.337 |

| **Structure** | **AAC(3)-IVa apoenzyme** | **AAC(3)-IVa^H154A^•APR** | **AAC(3)-IVa^H154A^•GEN** | **AAC(3)-IIb** | **AAC(3)-Xa** |
| --- | --- | --- | --- | --- | --- |
| **PDB code** | 6MN3 | 6MN4 | 6MN5 | 7LAO | 7LAP |
| **Data collection** |  |  |  |  |  |
| Space group | C2 | P2_1_2_1_2_1_ | P2_1_2_1_2_1_ | P2_1_2_1_2_1_ | P6_3_22 |
| Unit cell  *a*, *b, c* (Å)  α, β, γ, (°) | 114.2, 55.3, 94.3  90, 102.6, 90 | 77.6, 103.5, 264.9  90, 90, 90 | 77.6, 131.9, 266.9  90, 90, 90 | 43.2, 61.4, 112.0  90, 90, 90 | 161.5, 161.5, 138.7  90, 90, 120 |
| Resolution, Å | 30.00 – 2.39 | 30.00 – 2.80 | 40.0 – 2.58 | 40.0 – 1.92 | 50.0 – 2.04 |
| R*_merge_^a^*  R*_pim_* | 0.141 (0.997)  0.080 (0.572) | 0.183 (1.771)  0.063 (0.613) | 0.086 (0.542)  0.048 (0.371) | 0.080 (0.332)  0.037 (0.159) | 0.098 (1.074)  0.024 (0.396) |
| CC_1/2_ | ND | ND | 0.703 | 0.524 | 0.593 |
| *I* / σ(*I)* | 9.98 (1.25) | 12.77 (1.40) | 14.08 (1.08) | 26.19 (3.13) | 31.42 (1.08) |
| Completeness, % | 98.8 (99.9) | 95.2 (97.1) | 95.7 (82.0) | 95.8 (79.8) | 99.5 (92.9) |
| Redundancy | 3.9 (3.9) | 9.0 (8.9) | 3.7 (2.5) | 5.4 (4.7) | 17.2 (6.6) |
| **Refinement** |  |  |  |  |  |
| Resolution, Å | 30 – 2.39 | 29.33 – 2.80 | 38.4 – 2.58 | 35.32 – 1.92 | 49.39 – 2.04 |
| No. unique reflections:  working, test | 22587, 1129 | 63643, 3627 | 83240, 2000 | 22528, 2167 | 66736, 3279 |
| *R*-factor/free *R­*-factor^b^ | 18.2/22.8 (27.1/32.2) | 26.1/32.1 (35.3/41.1) | 18.7/22.1 (26.8/28.1) | 18.0/22.9 (23.6/29.4) | 16.5/19.8 (28.7/31.5) |
| No. refined atoms, molecules  Protein Aminoglycoside  Acetyl-CoA/CoA  Solvent  Water | 3921, 2  N/A  N/A  3  176 | 11564, 6  186, 5  N/A  5  330 | 11824, 6  186, 6  N/A  422  519 | 2045, 1  N/A  N/A  17  211 | 4421, 2  N/A  N/A  58  678 |
| *B*-factors  Protein  Aminoglycoside  Acetyl-CoA/CoA  Solvent  Water | 48.7  N/A  N/A  53.3  39.6 | 90.4  111.0  N/A  86.7  61.4 | 91.2  142.9  N/A  98.0  72.6 | 50.8  N/A  N/A  70.3  45.7 | 53.1  N/A  N/A  105.8  63.2 |
| r.m.s.d.  Bond lengths, Å  Bond angles, ° | 0.004  0.803 | 0.006  1.071 | 0.006  0.879 | 0.014  1.260 | 0.012  1.118 |

*values in brackets refer to highest resolution shells.

^a^*R*_merge_ = Σ_hkl_Σ_j_|*I*_hkl.j_ - 〈*I*_hkl_〉|/Σ_hkl_Σ_j_I_hk,j_, where *I*_hkl,j_ and 〈*I*_hkl_〉 are the *j*th and mean measurement of the intensity of reflection *j*.

^b^*R*_pim_ = Σ_hkl_√(n/n-1) Σ^n^_j=1_|*I*_hkl.j_ - 〈*I*_hkl_〉|/Σ_hkl_Σ_j_I_hk,j_

^c^value refers to highest resolution shell.

^d^*R* = Σ|F_p_^obs^ – F_p_^calc^|/ΣF_p_^obs^, where F_p_^obs^ and F_p_^calc^ are the observed and calculated structure factor amplitudes, respectively.

ND = not determined
